# Cryptic phylogeographic history sheds light on the generation of species diversity in sky-island mountains

**DOI:** 10.1101/199786

**Authors:** Kai He, Tao Wan, Klaus-Peter Koepfli, Wei Jin, Shao-Ying Liu, Xue-Long Jiang

## Abstract

Biodiversity hotspots should be given high priority for conservation under the situation of global climate change. The sky islands in southwestern China are characterized by extraordinarily high species diversity and are among one of the world’s top biodiversity hotspots. However, neither the actual species diversity in this region or mechanisms generating this diversity are well explored. Here, we report on the phylogeographic analysis of the long-tailed mole (*Scaptonyx fusicaudus*), a semi-fossorial mammal that inhabits the montane cool forests across the Chinese sky islands and is considered to represent one species divided into two subspecies. Analyses using DNA sequence data from one mitochondrial and six nuclear genes revealed that populations inhabiting different mountains exhibited exceptionally strong geographic structure. The lowlands and large rivers act as “soft” and “hard” barriers to dispersal, respectively, isolating evolutionary lineages for up to 11 million years. Our results suggest that the mountain ranges act as interglacial refugia buffering populations from climate fluctuations, further facilitating allopatric diversification. Strikingly, species delimitation analyses suggests that the long-tailed mole may comprise 18 operational taxonomic units and 17 putative species. Our results suggest that for low-vagility species, the complex topography of the Chinese sky islands has shaped genetic diversity and structure and promoted exceptional diversification through a combination of eco-environmental stability as well as geographic fragmentation. The patterns observed in *S. fusicaudus* may be representative for other cold-adapted species, reflecting the generation of mammalian faunal diversity in the sky-island mountains of southwestern China.

## INTRODUCTION

As the climate continues to change, one in six species are facing an accelerating risk of extinction [1]. This is especially true for narrowly endemic species [2]. To reduce the impact of climate change,optimizing the spatial priorities of conservation and protecting such areas from further anthropogenic activities could be a practical strategy [3]. Biodiversity hotspots should have the highest priority for conservation, because these areas represent only 2.3% of Earth’s land surface, but harbor more than half of the world’s plant species as endemics, as well as nearly 43% of birds, mammals, reptiles and amphibians [4].

Many of the biodiversity hotspots are low-latitude mountains that occur on different continents such as the mountains of the Eastern Afromontane in Africa, the Madrean pine-oak woodlands in North and Central America, the tropical Andes in South America, and the Himalayas as well as the mountains of southwestern China in Asia [4]. The extraordinarily high species diversity in these mountains are supported by various climatic and environmental conditions, resulting in altitudinal zonation and biologically complex ecosystems [5]. In addition, most low-latitude mountain regions are characterized by complex geographic features, composed of a series of discrete mountain archipelagos, also known as “sky islands” [6]. In a sky-island complex, each mountain is a fragmented habitat, surrounded by lowlands, which act as geographic barriers to montane species, leading to “speciation via isolation” in allopatry or parapatry [7]. The sky islands also act as refugia. Species can find hospitable habitat by moving up and down a few hundred meters and thereby survive climate fluctuations and avoid extinction. Thus, a large number of ancient relict species can be found on sky islands, known as the “species museum” effect [8]. Climatic changes may result in periodic distributional shifts of montane species, promoting allopatric speciation. Consequently, sky islands may also act as “species pumps”.

The mountains of southwestern China (also known as the Hengduan Mountains) and the adjacent mountains are among one of the world’s top global biodiversity hotspots [4]. The mountains span approximately 10° of latitude (N24°-N34°) and were recently recognized as a sky-island complex [9, 10]. This area is similar to other sky islands on Earth in its extremely complex topography, but distinctive by its well-developed river systems. The river canyons embedded between mountain ridges are comparable to the largest river basin in the world [11]. The sky islands in southwestern China are relatively young compared with the other sky islands in the world. The uplift started before 23 million years ago (Ma), but the major surfaces uplifted after 13 Ma [12]. Nevertheless, approximately 40% of the vascular plant and 50% of the mammal species in China are distributed in this area, a large portion of which are endemic. Many ancient and relic species are distributed in these mountains, including the giant panda (Ursidae),red panda (Ailuridae), shrew-like moles (Talpidae*, Uropsilus*), and gymnures (Erinaceidae*, Neotetracus*),demonstrating the buffering capacity of these mountains against climate change over geological timescales.

Despite the recognition as a biodiversity hotspot, the actual species diversity within the Hengduan Mountains or the mechanisms behind its generation is not well understood. Recent studies have shown that species diversity in this area is highly underestimated [13], and recently, the area has become a significant hotspot for the discovery of new species [e.g., 14, 15]. Inter- and intraspecific phylogeographic studies have revealed strong geographic structuring of lineages, which were associated with the complex topography of the region, geomorphologic changes (i.e., uplift of the mountains), or Pleistocene glaciations [16, 17]. However, the relative degree to which they promote diversification and speciation is not well known [10]. For example, previous studies showed the deep river valleys such as the Dadu, Min,and Yalong Rivers in the north sub-region as well as the Mekong and Salween Rivers in the southern sub-region of the sky islands (see **Fig. 1**) could act as barriers to dispersal of terrestrial small mammals resulting in intraspecific diversification [17, 18], while it is not known when these barriers appeared. Moreover, the montane archipelagos to the east and southeast of the Sichuan Basin including Mts. Bashan and Dalou in between of Mts. Jiaozi and Qinling (Fig. 1) have received little attention [10]. Although these archipelagos are quite discrete, they are inhabited by cold-adapted species, and might act as cryptic refugia and corridors of dispersal for montane species [17, 19].

**Figure 1.**
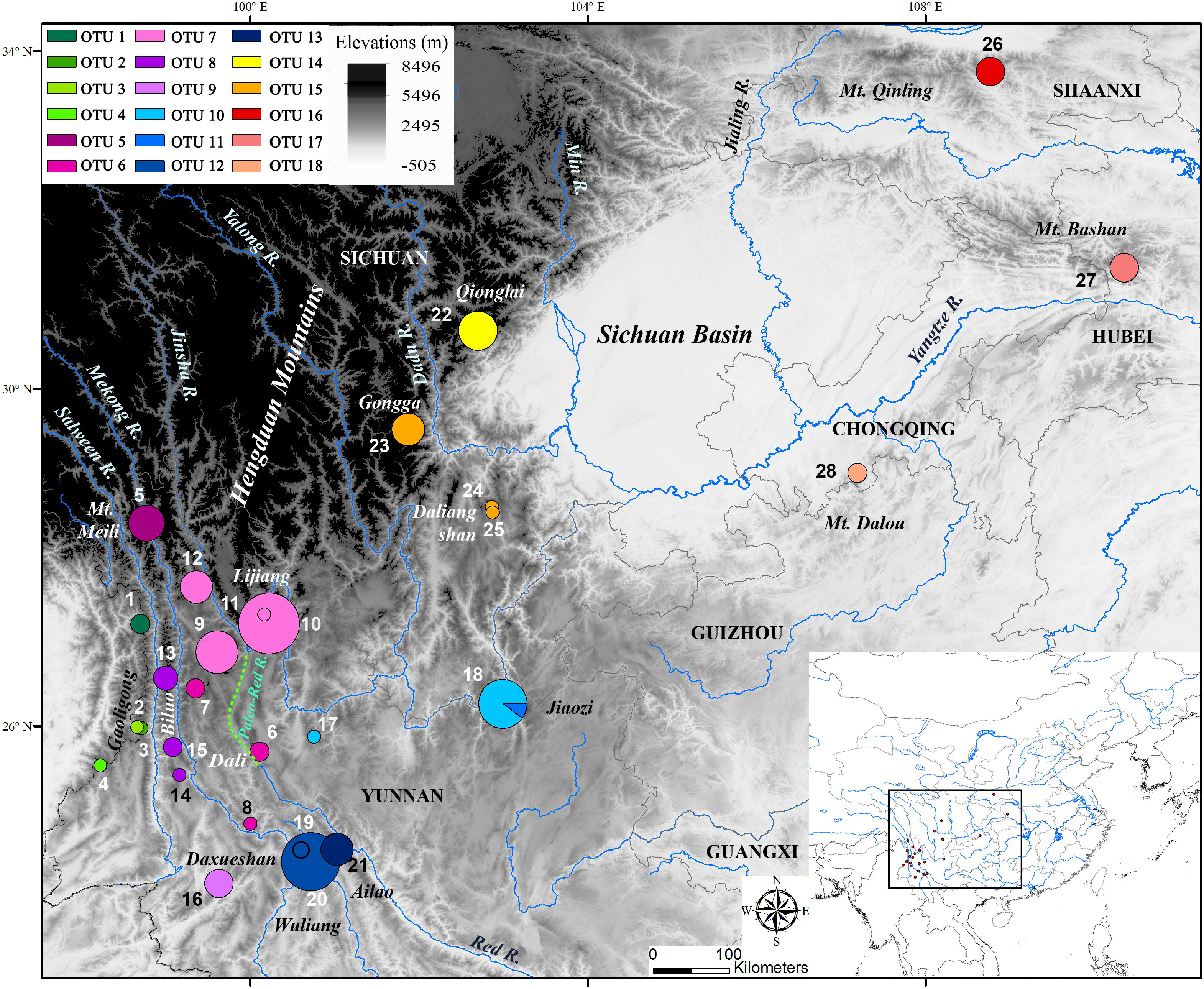
A map showing the sky islands in southwestern China, and the localities of *S. fusicaudus* samples collected for the current study. The locality numbers refer to those in Material S1.

The long-tailed mole (*Scaptonyx fusicaudu*s) is the sole representative of a monotypic genus within the family Talpidae (moles, shrew moles, desmans, and their allies), and is divided into two subspecies [20]. It is a small (weight 8-14 grams) semi-fossorial mammal mostly distributed in southwestern China. It is considered a relic species because its closest relatives are distributed in Japan (*Dymecodon* and *Urotrichus*) and North America (*Neurotrichus*), which diverged from the long-tailed mole approximately 30 million years ago [13]. It is also characterized by a strikingly deep intraspecific divergence (~16million years) in our previous estimation, which is more ancient than the most recent common ancestor(MRCA) of the shrew-like moles (*Uropsilus*), and that of the fossorial mole tribe Talpini [13]. We consider this species as an ideal model for phylogeographic study for the following reasons: i) it is a low-vagile species exclusively distributed at middle to high elevations. The populations are physically isolated among discrete mountain islands, so that its evolution could be strongly shaped by endemic geographic features as well as geomorphologic history; ii) it only inhabits humid environment, so that its genetic diversity could be shaped by climate change and following habitat turnover; iii) it is widely distributed through the sky-island mountains including those surrounding the Sichuan Basin, so it is good representative to reveal potential different effect in each sub-region or along the latitude gradient; and finally iv) its evolutionary time within these mountains is long enough to be comparable with the uplift history of the mountains, so that its evolution could be associated with the generation of species diversity in the sky islands.

In the present study, we analyzed mitochondrial and nuclear DNA sequences of long-tailed moles collected from throughout the sky islands of southwestern China to analyze genetic structure, estimate evolutionary relationships, and conduct species delimitation, approximate Bayesian computation and niche modeling analyses. We aimed to recover the phylogeographic patterns and the timescale for diversification, and unravel the underestimated species diversity. Furthermore, we examine to what degree geographic versus climatic factors have preserved and shaped the diversity found in long-tailed moles and thereby shed light on the underlying mechanisms that have driven diversification and speciation in the sky islands of southwestern China.

## MATERIALS AND METHODS

### Study area and sampling strategy

Our study areas include the northern Hengduan mountains (also known as the western Sichuan mountains; Nos. 22-25 in Fig. 1), the south Hengduan mountains (northwestern Yunnan; Nos. 1-7, 9-15, 17), the southern outliers of the Hengduan Mountains (nos. 8, 16, 19-21), montane archipelagos distributed from the south to the northeast of the Sichuan Basin including Mts. Jiaozi (no. 18), Dalou (no. 28), Bashan (27) and Qinling (no. 26). The study areas cover nearly all known distribution range except in adjacent northern Myanmar and northmost Vietnam. The sample localities were isolated by Dadu, Jinsha, Mekong, Red, and Salween Rivers, all of which likely act as barriers to dispersal for small mammals. The western Sichuan Mountains are less patchy than the other areas. The northwestern Yunnan is characterized by the most complex topography because the mountain chains are isolated by the Jinshan, Mekong and Salween Rivers, and also known as the Longitudinal Range-Gorge Region. The river valleys are as low as 300 m above sea level, supporting only Savanna vegetation and are unlikely suitable for the long-tailed moles. We collected 113 specimens of long-tailed moles from 1997 to 2013 (**Electronic supplementary material, Table S1**), four of which were sequenced in a previous study [13]. Skin and skull specimens and tissue samples were deposited in the Kunming Institute of Zoology (KIZ), Chinese Academy of Sciences (CAS) and the Sichuan Academy of Forestry (SAF).

### Sequence acquisitions

We extracted total DNA using the standard phenol/chloroform protocol [21]. Sample localities are shown in **Table S1**. We amplified one complete mitochondrial gene (*CYTB*) and six nuclear gene segments including *ADORA3*, *APP, ATP7*, *BCHE*, *BRCA1*, and *RAG1* using primers provided in **Table S2 (Electronic supplementary material)**. Sequences were assembled and edited using Lasergene v7.1 (DNAStar, Madison, WI) and aligned using MUSCLE (Edgar, 2004). Sequences of other shrew moles including *Dymecodon pilirostris* (n=1), *Neurotrichus gibbsii* (3) and *Urotrichus talpoides* (1) were obtained from GenBank (**Electronic supplementary material, Table S3**), and *Condylura cristata* (1) was used to root the trees. From the samples we collected, we obtained 113 *CYTB* (1140 bp), 112 *ADROA3* (358 bp), 111 *ATP7* (666 bp), 112 *APP* (640 bp), 103 *BCHE*, 113 *BRCA1* (969 bp), and 104 *RAG1* (1047 bp) sequences. Twenty-three sequences (3%) are missing in our alignments.

### Mitochondrial gene tree, tree-based and distance-based species delimitations

We estimated the *CYTB* gene tree, which was also used for tree-based species delimitation. We partitioned the *CYTB* gene by codon position and evaluated the best-fitting partitioning scheme and evolutionary models (**Electronic supplementary material, Table S4**) using PartitionFinder v1.1 [22]. Bayesian analyses were performed using BEAST v1.8 [23]. We constrained the monophyly of Dymecodon + *Neurotrichus* + *Scaptonyx* + *Urotrichus* based on our best knowledge regarding their relationships [13]. For each analysis, we selected a lognormal relaxed-clock model, a birth–death tree prior, and ran the Markov chain Monte Carlo for 50 million generations, with a sampling interval of 5,000 generations. The analyses were repeated four times and were performed on the CIPRES Science Gateway [24]. Posterior distributions and effective sample sizes were calculated using Tracer v1.6.

We used two approaches including Poisson Tree Processes (PTP) and the Generalized Mixed Yule Coalescent (GMYC) to conduct tree-based species delimitation. The Bayesian tree with branch lengths representing substitutions/site was used for the PTP analyses and implemented in the bPTP server [25]. We used the default settings for all parameters and confirmed the convergence by examining the trace plot. The tree with branch lengths proportional to time was used for GMYC analyses and implemented using the R package "splits” (http://r-forge.r-project.org/projects/splits). For the distance-based species delimitation, we calculated the number of operational taxonomic units (OTUs) using USEARCH v8.1 [26]. We used a minimum identity of 97% as the “radius” of an OTU.

### Nuclear gene trees and molecular dating

We estimated the concatenated nuclear gene tree using phased alleles. We phased each of the nuclear genes of the ingroup taxa using PHASE [27] and implemented in DnaSP v 5.1 [28]. The best-fitting partitioning schemes and evolutionary models were estimated for each gene using PartitionFinder v1.1. To minimize the effect due to missing data, we only included samples from which all six genes were successfully sequenced. This dataset included 198 phased alleles representing 99 samples of *S. fusicaudus* plus six unphased outgroup samples as mentioned above.

We estimated the Bayesian trees as described above. The phased gene alignments were concatenated, and defined by genes and codon positions, and used for estimating the best partitioning scheme and evolutionary models with PartitionFinder (**Electronic supplementary material, Table S4**). Two fossil-based calibrations were selected for divergence time estimation. The oldest known fossil of shrew moles (*Oreotalpa*) from the strata of the latest Eocene [29] was used to calibrate the root of the tree, and the oldest known *Scaptonyx* fossil specimens from the early Pleistocene strata [30] from Shennongjia (Mt. Bashan), Hubei, was used to calibrate the stem group of the population. The justifications for placement of these calibration priors are detailed in **Text S1 (Electronic supplementary material)**.

The estimated MRCA of *S. fusicaudus* was strikingly ancient (the early Miocene, see Result Section), with a high level of uncertainty (i.e., large 95% confidence intervals). To increase accuracy, we compared four competing hypotheses of divergence times by constraining the root age of *S. fusicaudus* to be at i) the Oligocene/Miocene boundary (23.03 Ma; H0); ii) the Middle/Late Miocene boundary (11.61 Ma; H1); iii) the Miocene/Pliocene boundary (5.40 Ma; H2) and iv) the Pliocene/Pleistocene boundary (2.40 Ma; H3). These four hypotheses were compared using Bayes factors estimated with the path sampling (PS) and stepping-stone sampling (SS) methods [31] and implemented in BEAST v1.8. We employed normal distributions for the priors of the time of the MRCA for the root age and used 0.5 as the standard deviation (SD) in all four analyses. To calculate the Bayes factor the marginal likelihood of each analysis was estimated with 1,000,000 steps. The two fossil-based calibrations were used as “checkpoints”, constraining the times of diversifications. Similar approaches have been employed in previous studies [32, 33]. We considered the Bayes Factor of ≥ 10 as very strong evidence against the alternative hypothesis [34].

### Structure and coalescent species delimitation using nuclear genes

We used the phased nuclear genes to estimate population structure for *S. fusicaudus* using the correlated admixture model in STRUCTURE v2.3.4 [35]. Because the mitochondrial gene tree showed a strong geographical pattern, we assigned individuals from the same mountain area to their own populations. To reduce computational cost, we split the samples into three datasets, corresponding to the three major clades recovered in both the mitochondrial and nuclear gene trees. The first dataset included 10 phased samples from Mt. Gaoligong, and we tested K values from 1 to 4. The second dataset included 14 phased samples from the western Sichuan mountains, and we tested K values from 1 to 4. The third dataset included all the other samples (n=96), and we tested K values from 3 to 15. Each run consisted of 1,000,000 step Markov chain Monte Carlo (MCMC) replicates after a burn-in of 100,000 replicates. We repeated the analysis 20 times for each dataset. We used a “LargeKGreedy” algorithm for aligning multiple replicate analyses as implemented in CLUMPK using the default parameters [36, 37], and determined the optimal number of clusters (i.e., the best K value) for each dataset using ln(Pr(X|K) values (see STRUCTURE manual).

We conducted coalescent-based species delimitation analyses of the phased nuclear genes using BPP v3.1 [38] and BEAST v2.3 [39]. Because the algorithms of BPP were not sensitive to missing data, we included all ingroup samples. To avoid the potential of poor mixture of the reversible-jump MCMC algorithms, we split our samples into four datasets, corresponding to clade I, clade II, subclade A, and subclades B+C, according to the topology of the concatenated nuclear gene tree. Samples were assigned to candidate species based on the results of STRUCTURE. We followed Yang [40] for running the BPP pipelines. First, the genetic distances in each dataset were calculated using MEGA v7.0 [41], and the values were used for the priors of “thetaprior." We then estimated the priors for “tauprior” using the A00 model. The species tree topologies were fixed according to the mitochondrial gene tree for parameter estimations (A00). To avoid false positive errors due to incorrect guide trees [42], we used the A11 model to allow simultaneous species delimitation and species-tree estimation [43]. To allow variable mutation rates among loci, we set the parameter α to 20 for “locusrate." We used both algorithms 0 and 1, and repeated the analyses for each dataset at least six times with combinations of different parameters and priors. For each run, 5,000 samples were collected by sampling every 10 iterations after a burn-in of 5000 iterations. We used a conservative criterion, recognizing posterior probabilities ≥ 0.95 as strong support for putative species.

We also conducted species delimitation and species tree estimation using the BEAST2 package STACEY v1.1, which is also based on a multispecies coalescent model [44]. To reduce computational costs, we assigned an HKY+G model to each of the nuclear genes and employed a strict molecular clock model with a mean evolutionary rate of 0.0015 based on the results of the BEAST analyses using the concatenated nuclear gene dataset. Samples were assigned to candidate species based on the results of STRUCTURE. The analyses were conducted using BEAST2 v2.3.3.

#### Approximate Bayesian Computation (ABC)

DIYABC v.2.1 [45] was used for ABC-based scenario comparisons, and to examine potential gene flow along the mountains to the east of the Sichuan Basin, which may serve as stepping stones among isolated populations. To reduce computational cost, a subset of samples (n=40) from Mts. Ailao, Bashan, Dalou, Jiaozi, Wuliang and Qinling (i.e., subclades A and C except for the single individual from Mt. Wuding) were selected and were assigned to six populations according to their distributions. All six nuclear genes were included in one group of the mutation model. We selected HKY as the substitution model and estimated the percentage of invariant sites and the shape of the gamma distribution using MEGA6 [46].

We performed four runs of analyses to test competing scenarios. The priors for divergence times, population sizes, and mutation rates are given in **Text S2 (Electronic supplementary material)**. First, 10 we compared four diversification scenarios to test for the potential of simultaneous divergence. Second, we compared the best topology with three gene-flow scenarios. Third, we compared four alternative scenarios regarding the time of gene flow. Finally, we compared scenarios regarding demographic changes (**Test S2**). We simulated all summary statistics within populations except for private segregating sites, and all summary statistics between two populations. We simulated a minimum of 10,000 datasets for each comparison. After simulations, we ran pre-evaluations to confirm that the simulated datasets were not significantly different from the observed data. Then we compared scenarios based on a logistic regression estimation of the posterior probabilities (PP) of each scenario. The best scenario was selected based on non-overlapping 95% confidence intervals (CIs) of PP. Two scenarios with overlapping 95% CIs were not considered as significantly different and the simpler scenario was then selected.

#### Species distribution modeling (SDM)

We obtained 76 geographic coordinates in our field work and adopted another two reliable coordinates from previous studies [47, 48]. To reduce the effect of sample bias, we consequently subsampled 63 coordinates by retaining one record per grid cell of 0.2° × 0.2° using the R package Raster. Nineteen climate variables for the present day were downloaded from the WorldClim website. We extracted the 19 variables for each coordinate using DIVA-GIS v7.5. Nine variables that were not strongly correlated (Pearson’s R < 0.9; Bio 1, 2, 3, 5, 7, 12-15) were retained for further analyses. We conducted principal components analyses (PCA) and discriminant factor analyses (DFA). We assigned all samples into five groups based on their phylogenetic positions (clades and subclades) in the DFA. We used MaxEnt v.3.3.3k to conduct niche modeling [49], calculating the potentially habitable areas in the present day, during the last interglacial maximum (LIGM), and during the last glacial maximum (LGM). For estimating niches in LGM and LIGM, climatic datasets were also downloaded from WorldClim. The area under the receiver operating characteristic curve (AUC) was used to evaluate the accuracy of our models, and AUCs higher than 0.9 were considered as an indicator of good projecting.

## RESULT

### Structure and diversity of the mitochondrial data

Three major clades were recovered in the *CYTB* gene tree (Clades I, II and III; **Fig. 1**), showing a high level of geographic structure. Five specimens from Mt. Gaoligong (the westernmost mountain in Yunnan) formed a basal position of *S. fusicaudus* (clade I). The samples from the western Sichuan mountains composed the second major clade (clade II). Clade III was comprised of three subclades including i) samples from Mt. Qinling (Shaanxi), Mt. Bashan (Hubei) and Mt. Dalou (Chongqing) (subclade A); ii) samples approximately distributed to the west of 100.5°E longitude in Yunnan, but to the east of the Mekong River (subclade B); and iii) samples from middle and eastern Yunnan, approximately to the east of the 100.5°E longitude (subclade C). The mitochondrial *CYTB* gene was characterized by high haplotype and nucleotide diversities (Hd = 0.99, Pi = 0.09; **Electronic supplementary material**, **Table S5**).

### Nuclear gene tree and ancient divergence times

In the concatenated nuclear gene tree, three major clades (I, II and III) were recovered (**Fig. 2**), and the monophyly of 15 mitochondrial lineages was strongly supported. Within clade III, the monophyly of subclade A was highly supported (PP=0.99). Lineages within subclades B and C of the *CYTB* gene tree showed an alternative pattern of relationships: the population from Mt. Jiaozi formed a basal clade to the other populations (PP=0.97), the single specimen from Mt. Wuding was recovered as a distinct lineage, and relationships among populations from Dali and Lijiang mountain ranges were not resolved.

**Figure 2.**
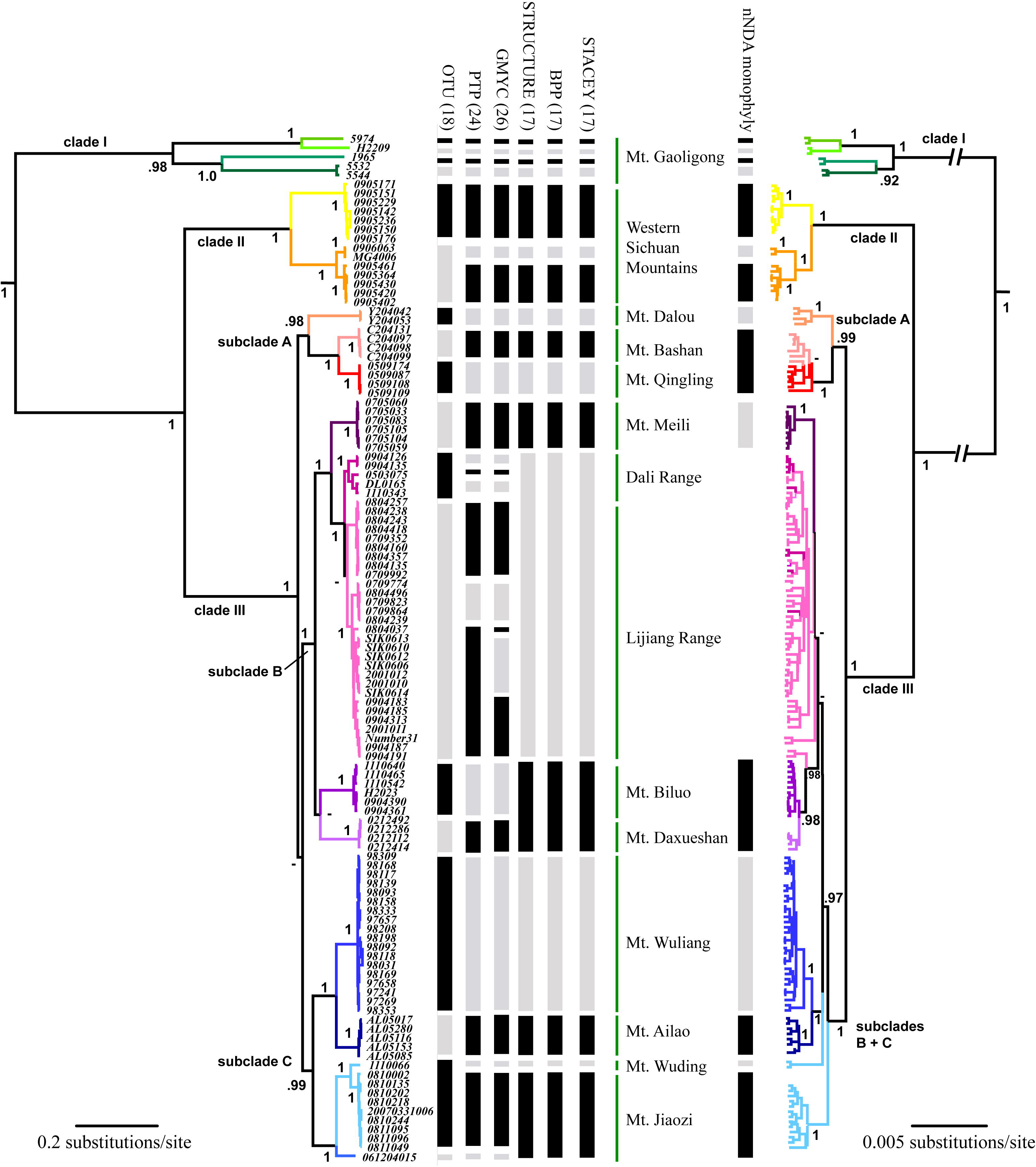
A comparison of the mitochondrial gene tree (left) and the concatenated nuclear gene tree (right). The estimated OTUs, the result of clustering analysis using STRUCTURE, and the results of species delimitation analyses using bPTP, split, BPP and STACEY are shown in the middle. The numbers at the nodes refer to Bayesian posterior probabilities. Branch lengths represent substitutions per site (scale bars shown).

Divergence time estimation based on nuclear genes supported the tMRCA of *S. fusicaudus* at 25.04 Ma, but this was associated with a large uncertainty (95% confident interval [CI] = 37.68-15.29; **Fig. S1*A*, Electronic supplementary material**). The age was three times older than that between the two shrew mole genera in Japan (*Dymecodon* and *Urotrichus*; 6.65 [95% CI=11.12-3.38]). The Bayesian-based scenario comparison (**Table 1**) rejected hypotheses H2 and H3 (Δ PS and Δ SS ≥ 15), but could not reject H1 (Δ PS and Δ SS ≤ 2; **Fig. 3*A***), indicating the tMRCA is no younger than 11 Ma (**Fig. S1*B-E***).

**Table 1.** Model comparison for diversification scenarios. Divergence times and Bayes factors estimated with the path and stepping-stone sampling (PS/SS) methods are presented.

| Hypothesis | TMRCAs of *S. fusicaudus* | Prior /Posterior of TMRCAs (Ma) | path sampling | $\Delta$ PS | stepping-stone | $\Delta$ SS | *Dymecodon* / *Urotrichus* (95%CI) |
| --- | --- | --- | --- | --- | --- | --- | --- |
| H0 | Oligocene / Miocene | 23.03 / 23.05 | **-15672** | - | -15676 | 2 | 6.12 (9.03-3.48) |
| H1 | middle / late Miocene | 11.61 / 11.84 | -15673 | 1 | **-15674** | - | 4.84 (8.70-1.91) |
| H2 | Miocene / Pliocene | 5.4 / 6.08 | -15688 | 16 | -15689 | 15 | 3.32 (7.09-1.11) |
| H3 | Pliocene / Pleistocene | 2.4 / 3.83 | -15701 | 29 | -15702 | 28 | 2.50 (5.57-0.71) |

**Figure 3.**
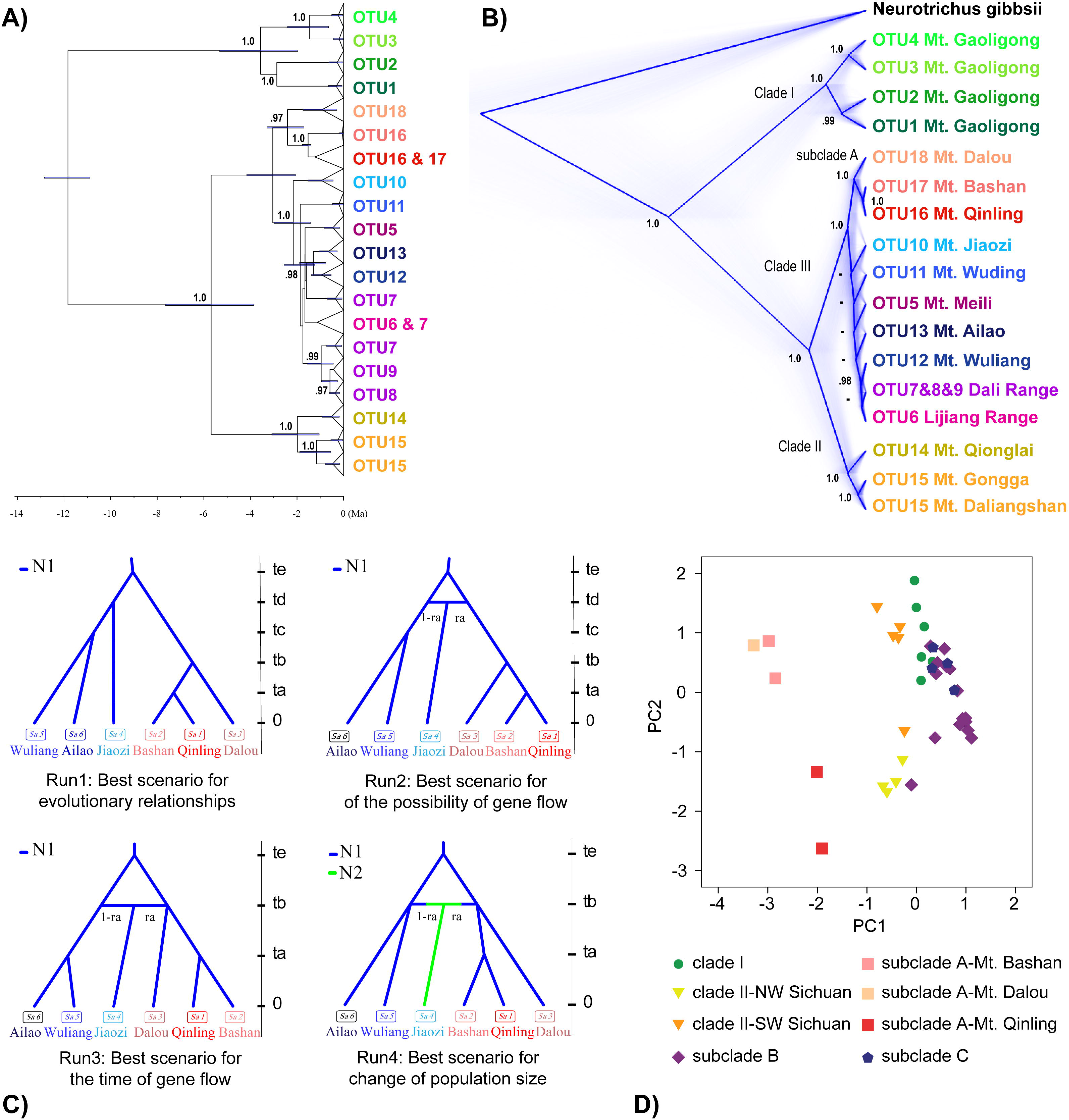
Divergence times estimated using concatenated nuclear genes (A), and a coalescent species tree estimated using the STACEY model in BEAST2 (B). Values at nodes represent posterior probabilities (PPs). PPs <0.95 are not shown. Horizontal bars in (A) represent 95% confidence intervals (CIs) for divergence times. The timetree shown is the one estimated using the H1 scenario for calibration priors, in which the MRCA was constrained at 11.61 Ma. (C) The best-fitting scenarios selected in four runs of the ABC analyses. (D) Principal components analyses (PCA) for the habitats of clades I and II and subclades A-C. Plots of scores on PC1 and PC2 were from PCA of nine climatic variables.

### Genetic structure, species delimitation and species tree

When using 97% similarity as a criterion, the USEARCH analysis recovered 18 OTUs of *S. fusicaudus* (**Table 1**, **Fig. 2**). Mitochondrial tree-based analyses including PTP and GMYC recognized 24 (confidential interval [CI] = 22 - 30) and 26 (CI=23-29) putative species, respectively (**Fig. 2**). STRUCTURE, as well as BPP and STACEY delimitation analyses, supported 17 isolated populations or putative species. The coalescent species tree recovered using the STACEY model was similar to that of the concatenated nuclear gene tree, although relationships among the putative species recovered in subclades B + C in the mitochondrial gene tree were remained unresolved (**Fig. 3*B***).

### Detection of gene flow using DIYABC

DIYABC analyses were limited to the populations to the east (Nos. 26, 27 in **Fig. 1**), north (No. 28) and south (Nos. 18-21) of the Sichuan Basin to detect the possibility of long-distance dispersal among a set of discrete mountains (**Fig. 3C**). These analyses suggest that after the first split somewhere between the eastern (Mt. Dalou) and southern populations (Mts. Wuliang + Ailao), gene flow occurred bewteen these populations, resulting in admixture (admixture rate = 0.47 [CI=0.28-0.66]) at Mt. Jiaozi (No. 18 in **Fig. 1**) in northeastern Yunnan. The dispersal and admixture was accompanied by a demographic expansion (N2=4.36 *10^4^ [CI=3.17-5.85] > N1=2.91*10^4^[CI=2.35-3.57]).

### Species distribution modeling

In the PCA plot based on nine climatic variables (**Fig. 3D**), subclade A (comprising samples to the east and northeast of the Sichuan Basin) was well separated from the others along PC1. Clade I and subclades B and C plot together and partly overlap with each other, suggesting all populations from Yunnan inhabit similar environments, despite their distinct phylogenetic histories. A similar pattern was also revealed in the DFA (**Fig. S2, Electronic supplementary material**).

The results of SDM showed that suitable habitats in the present day are larger than during the Last Interglacial Maximum, but smaller than during the Last Glacial Maximum (**Fig. 4*A-C***). The SDM did not support the existence of suitable habitats during the Last Interglacial Maximum at Mt. Qinling and to the east of the Sichuan Basin. On the other hand, suitable habitats were always present in northwestern and northeastern Yunnan as well as in the adjacent southern Sichuan mountain areas during all three time periods (**Fig. 4*D***).

**Figure 4.**
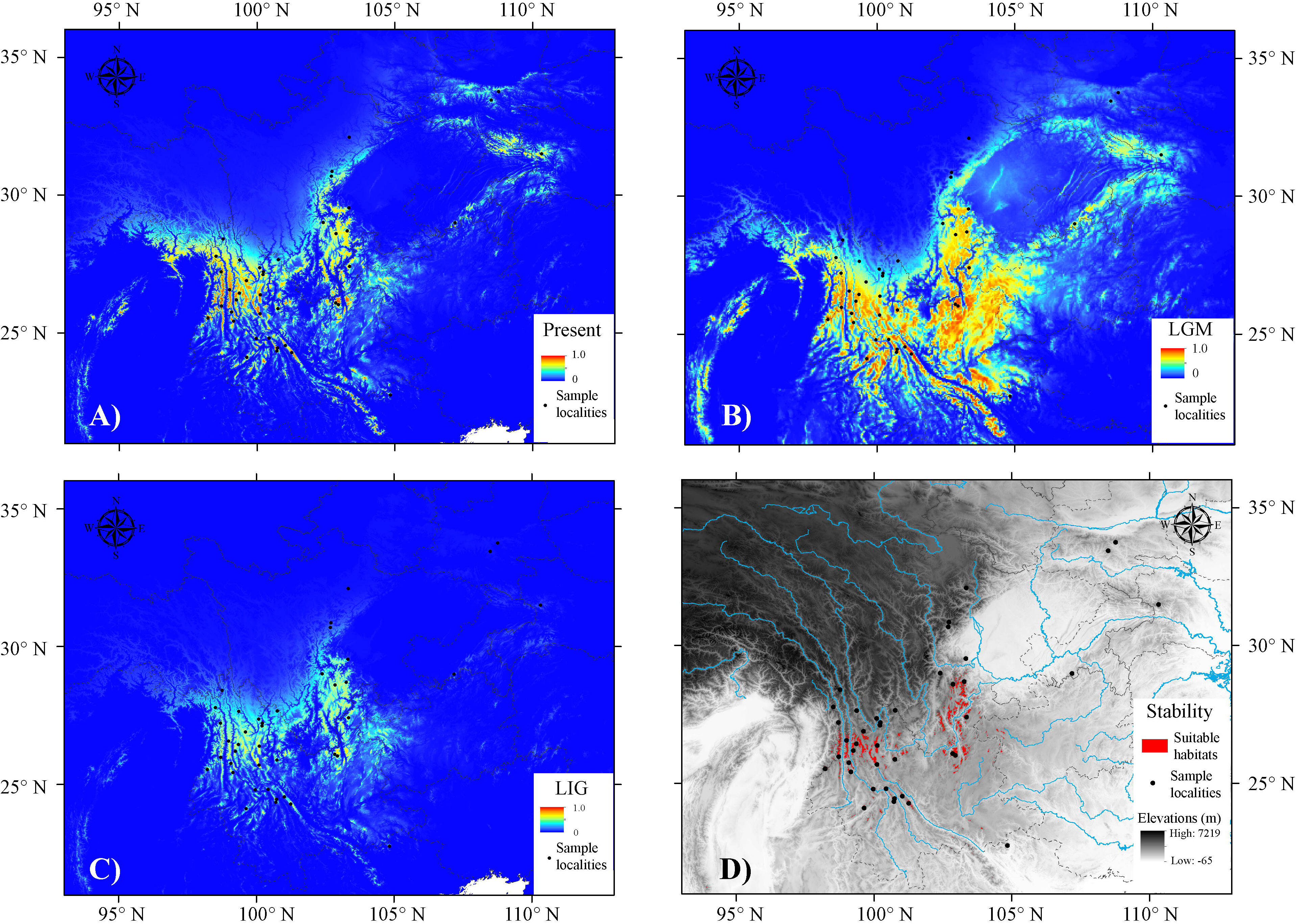
Ecological niche model projections for the distribution of *S. fusicaudus* in the present day (A), during the Last Glacial Maximum (B), and the Last Interglacial Maximum (C). Potential distributions with probabilities higher than 0.5 in the present day, the LGM and LIGM are extracted and shown in (D). **TABLES**

## DISCUSSION

Our results indicate that *S. fusicaudus* is characterized by strikingly ancient and *in situ* diversification of multiple lineages and high genetic diversity, which is unevenly distributed across the sky islands in southwestern China. The patterns we have uncovered represent, to the best of our knowledge, the most extreme case of geographic diversification in this area, and supports the idea that geographical factors could have played a more important role than climatic changes in shaping genetic diversity and geographic structure within the sky islands of southwestern China. Our results suggest that the mountains have acted as a buffer against climate change, and have provided continuously suitable habitats for *S. fusicaudus* since the early Late Miocene, which supports the hypothesis of the sky islands as a “museum” of ancient species and lineages. Our phylogeographic results suggest that the local rivers could act as “hard” barriers preventing gene flow, while the montane archipelagos act as stepping stones to facilitate dispersal.

### Striking diversification and underestimated diversity

The most striking finding of our study are the multiple ancient intra-specific splits among lineages within *S. fusicaudus*. The estimated ages on the basis of two fossil calibrations are characterized by wide confidence intervals, which is probably due to the uncertainty of the priors used to calibrate the timetree. Nonetheless, PS/SS-based scenario comparisons support the MRCA of *S. fusicaudus* to have split no younger than the earliest late Miocene, ~11 million years ago (**Figs. 3 and S1**), twice as old as the ancestor of the two shrew mole genera *Dymecodon* and *Urotrichus* (**Table 1**).

The ancient diversification of lineages is accompanied by a minimum of 17 and as many as 26 OTUs or putative species, as recovered in all mitochondrial-based and nuclear-based structure and species delimitation analyses (**Fig. 1**). A recent study showed that in a sky-island system, cryptic speciation as well as extreme population genetic structuring (but not speciation) could occur and the latter also result in oversplitting of lineages [50]. Distinguishing the two requires further taxonomic revision, and the patterns we found should be considered as hypotheses that requires validation based on morphological and ecological analyses. We did not perform morphological analyses due to the small sample size of skulls; in addition, most specimens have broken skulls. Nevertheless, the large number of OTUs and putative species recovered have important implications for conservation. Because each of them inhabit restricted mountain ranges (see below), loss of habitats could result in the extirpation of an entire distinctive genetic lineage or evolutionary significant unit (Ryder, 1986).

### Allopatric diversification shaped by the drainage and mountain systems

The second striking finding was the strong geographic structure of populations among different mountain ranges of the sky islands. Because *S. fusicaudus* is a montane inhabitant, and because the landscape in southwestern China is patchy, the habitats of *S. fusicaudus* are discontinuous as revealed in the SDM results (**Fig. 4*A***), which could have led to geographic isolation and genetic drift. As a result, each of the 18 mtDNA OTUs (most of which were also recovered by the nuclear genes) is endemic to specific mountain range (**Fig. 1**).

The deep rivers may be considered as "hard” barriers because they strengthen the barriers to dispersal. For example, clades I and III diverged in the Late Miocene but geographically they are only isolated by the Salween River (**Fig. 1**). PCA and DFA using climatic variables of the present day suggests habitats on both sides of the Salween River were similar, at least much more similar than between Yunnan and Shaanxi, which are inhabited by subclades A-C (**Figs. 3D & S2**). Therefore, the 11-million-year divergence between clades I and II is due to geographic isolation instead of adaptation to a new environmental habitat. The barrier effect could be due to: i) the Salween River, which acts as a physical barrier to dispersal, ii) the extremely hot and dry climate due to the foehn winds in the valley (see He and Jiang 2014), and iii) the very limited dispersal ability of the species. Similarly, the other large rivers including the Dadu, Mekong, and Red Rivers also seem to have acted as barriers, similar to the patterns revealed in previous studies [17, 18].

Basins and lowlands could be considered as “soft” barriers, allowing dispersal and gene flow during climate fluctuations [altitudinal movements; 51]. Conflicting gene trees and species delimitation patterns using mitochondrial and nuclear DNA suggest a recent mitochondrial introgression from Mt. Jiaozi to Mt. Wuding (Nos. 17 and 18 in **Fig. 1; Fig. 2**). Similarly, the close genetic affinity between the populations in Mt. Qinling and eastern Yunnan (clade III) was recovered in all phylogenetic analyses (**Figs. 2, 3A-B**) as well as the ABC results supporting a gene flow along the mountains to the east of the Sichuan Basin. The results suggest these montane archipelagos have acted as a stepping-stone corridor, facilitating migration of *S. fusicaudus* between montane habitats. This result was *unexpected*, because the Hengduan Mountains throughout western Sichuan have been hypothesized to play this role [52], acting as the primary corridor between the Palearctic and Oriental realms [53, 54]. Compared with the Hengduan Mountains, the geomorphology of mountains to the east of the Sichuan Basin is patchier and lower in elevation. Importantly, the discovery of this cryptic corridor demonstrates montane species could be distributed around the Sichuan Basin in a ring, which is a precondition of the existence of ring species [55].

### Interglacial refugia maintaining genetic diversity

The geomorphology of the sky islands is likely the primary driver of diversification in *S. fusicaudus*, but it does not necessarily support diversity for extended periods of time. Instead, diversity is maintained over time in stable habitats during interglacial periods (namely, interglacial refugia).

According to the SDM results, the potentially suitable habitats of *S. fusicaudus* enlarged during the LGM and reduced during the LIGM (**Fig. 4 *B-C***). This is a general pattern for low-altitude montane inhabitants [56] and the term of refugia should be more specifically referred as “interglacial refugia” instead of “glacial refugia” [51]. Our phylogeographic and divergence time results suggest that suitable habitats for *S. fusicaudus* were available during most of the Pleistocene and beyond. This pattern is not unexpected because the mountains in this area have a wide altitudinal range-span, therefore montane inhabitants only need to migrate a few hundred meters in altitude to avoid extirpation.

The SDM analyses also supported more stable habitats in the southern distribution area (**Fig. 4*D***). We note that our SDM analyses were conducted only for the last 130,000 years, which is much shorter than the evolutionary timeframe estimated for *S. fusicaudus*. We also ignored any geomorphological changes due to the uplift of the Himalaya and Hengduan mountains. Furthermore, the populations in the Mts. Qinling, Bashan, and Dalou could have adapted to new climatic environments, as observed in the PCA and DFA plots (**Figs. 3*D* & S2**, which could result in an underestimation of the potential distribution in the northeastern part of the range of *S. fusicaudus*. Nevertheless, the patterns revealed in the SDM coincide with the unevenly distributed genetic diversity. Specifically, thirteen out of the 18 OTUs are distributed in the south, whereas Mt. Gaoligong has supported *S. fusicaudus* for up to 11 million years, while only 5 are distributed in the north.

### Implication for endemic species diversity of the mountains

The phylogeographic pattern observed in *S. fusicaudus* is the most extreme case that we have ever seen for a single species inhabiting southwestern China. Although it is taxonomically considered a monotypic genus and species, the >10-million-year evolutionary timeframe and wide distribution makes the long-tailed mole a good representative of endemic montane organisms with low vagility, and likely reflect the generation of species diversity in the sky-island mountains. Indeed, the unevenly distributed genetic diversity in *S. fusicaudus* analogue the species richness pattern of vascular plants, of which higher species richness and endemism was observed in the southern sub-region [57].

First, the species diversity is probably underestimated. The cryptic diversity we observed in *S. fusicaudus*, was also found in most of the mammalian genera we have examined [e.g., 58, 59], indicating existence of unknown species [14]. Second, the extraordinary species diversity should thanks to the highly stable environments, which buffer dramatic global cooling and desiccating since the late Miocene and periodical climatic changes during the Pliocene and Pleistocene. This is especially true for Mt. Gaoligong, which has possibly harbored *S. fusicaudus* for as long as 11 million years. The stabilized climatic condition makes the mountains a species “museum”. Third, the complex topography of the sky-island mountains has facilitated allopatric speciation even over long evolutionary timescales, resulting in the appearance of narrowly endemic species. Deep rivers such as the Salween and Mekong Rivers may act as strong barriers to dispersal, which could further lead to high species diversity and the unique fauna found in northwestern Yunnan such as on Mts. Gaoligong and Biluo. The allopatric diversification pattern is likely applicable for most cold-adapted as well as hot-adapted terrestrial organisms, although for hot-adapted organisms, the mountains likely act as barriers and low lands acting as corridors, conversely [60].

## ACKNOWLEDGEMENTS

We thank Mr. Chang-Zhe Pu for sample collections. This work was supported by the Key Research Program of the Chinese Academy of Sciences (KJZD-EW-L07), the Yunnan Applied Basic Research Projects (2014FB176) and the National Natural Science Foundation of China (31301869).

## DATA ACCESSIBILITY

DNA sequences have been deposited in GenBank nucleotide database under the accession codes XXXXXX to XXXXXXX.

## AUTHOR CONTRIBUTIONS

K.H. and X.L.J conceived the idea for the study. All authors contributed to the data collection. K.H. and T.W analyzed the data. K.H. and K.P.K. wrote the manuscript.

## ELECTRONIC SUPPLEMENTARY MATERIAL

**Table S1.** Samples used for the molecular work in the present study.

**Table S2.** Primers used for amplification and sequencing.

**Table S3.** Species and sequences used as outgroups for divergence time analyses.

**Table S4.** Partitioning schemes and evolutionary models used for the phylogenetic analyses of the concatenated nuclear gene dataset.

**Table S5.** Genetic diversity of each gene segment used in the study.

**Text S1.** Justifications for the two fossil-based calibration priors used in divergence time estimation analyses.

**Text S2.** Parameters, priors, scenarios and posterior probabilities for the four runs of ABC analyses.

**Figure S1.** Chronograms estimated using the concatenated nuclear gene da. Divergence times were estimated based on two fossil-based calibrations (A), and with additional constraints on the MRCA of *S. fusicaudus* (B-E). Branch lengths represent time. Node bars indicate the 95% CI for the clade age. The result of H1 is also presented in **Fig. 3A**.

**Figure S2.** Discriminant function analysis (DFA) for the habitats of clades I and II and subclades A-C. Plots of scores on functions 1 and 2 were from DFA of nine climatic variables.

